# Greater loss of female embryos during human pregnancy: A novel mechanism

**DOI:** 10.1101/418186

**Authors:** John F Mulley

## Abstract

Given an equal sex ratio at conception, we can only explain the excess of human males at birth by greater loss of females during pregnancy. I propose that the bias against females during human development is the result of a greater degree of genetic and metabolic “differentness” between female embryos and maternal tissues than for similarly aged males, and that successful implantation and placentation represents a threshold dichotomy, where the acceptance threshold shifts depending on maternal condition, especially stress. Right and left ovaries are not equal, and neither are the eggs and follicular fluid that they produce, and I further hypothesise that during times of stress, the implantation threshold is shifted sufficiently to favour survival of females, most likely those originating from the right ovary, and that this, rather than simply a greater loss of males, explains at least some of the variability in the human sex ratio at birth.

## Introduction

### The choosy uterus

Pregnancy can in part be viewed as a conflict between mother and offspring.^[1]^ Selection acts on maternal genes to limit the supply of resources to developing offspring so as to maximise (or at least stabilise) maternal fitness, whereas fetal genes are selected to maximise growth.^[1, 2]^ These selfish offspring may seek to maximise their own growth at the expense of future offspring, whereas maternal investment may vary with age, with younger mothers less likely, and older mothers more likely, to sacrifice their well-being for the benefit of their offspring.^[3, 4]^ Imprinting of maternal genes therefore serves to control the growth and/or function of the placenta. There is also potential conflict between maternal and paternal alleles, as fathers have an interest in improved survival of (their) current offspring even at the expense of future offspring, which may have a different father. With respect to offspring sex ratios, the Trivers and Willard Hypothesis proposes that as females deviate from the “average” condition they should bias the production of one sex over the other, driven by improved likelihood of producing grandchildren,^[5]^ and logic suggests that this sex ratio manipulation should occur early to minimise wasted maternal investment. In humans, the sex ratio at birth is male-biased, with around 1,055 males born per 1,000 females in England and Wales between 1927 and 2007 (Figure 1). Since implantation represents the first major instance of fetal-maternal conflict, I hypothesise that it is at this point that a large component of variation in the human sex ratio arises, facilitated by sensing of embryo quality by the uterine lining (endometrium),^[6–9]^ most specifically the degree of “*differentness*” from the mother. Implantation therefore represents a threshold dichotomy,^[10]^ where passing the threshold results in successful implantation and initiation of placentation (and at least a chance of further development) and failing results in loss, but where the threshold itself can vary within and between women. The mammalian implantation process is thought to have evolved from endometrial inflammation – a natural reaction of maternal tissues to a foreign body.^[11, 12]^ Whilst the initial stages of implantation are pro-inflammatory, post-implantation embryonic development requires an anti-inflammatory endometrial state, and there must therefore be a point of acceptance by the maternal tissues either at initial contact, or soon after as the embryo “invades” the endometrium and as the process of placentation progresses.^[7]^ The first few weeks of gestation place relatively little demand upon the mother and therefore involve little maternal investment, and so embryo quality control should occur early, with the fetal-maternal interface (i.e. the interaction between fetal ligands and maternal receptors) representing the front line in the battle between invading trophoblast and defending maternal tissues. Indeed, discussion of early fetal-maternal interactions is full of references to embryonic “invasion” of the endometrium, and the literature is full of war-like references to “fighting lines”,^[13]^ “no man’s land”,^[14]^ and the embryo as a “deceitful and treacherous enemy”.^[15]^ A more appropriate comparison for the very earliest stages of implantation at or soon after the initiation of embryo-maternal contact may be an interview, where the embryo seeks to make a favourable impression.^[14]^ The evolution of deeper implantation in placental mammals facilitated more thorough vetting of offspring, and these deeper forms of implantation may have evolved to reduce the “ease” by which a mother may reject an embryos through sloughing of superficial layers of endometrium.^[14]^ This process, together with rapid evolution of placental proteins^[3]^ reflects the fetal-maternal arms race. How might this maternal vetting of embryos occur, and why might it preferentially target female embryos to result in a male-biased sex ratio at birth?

**Figure 1.**
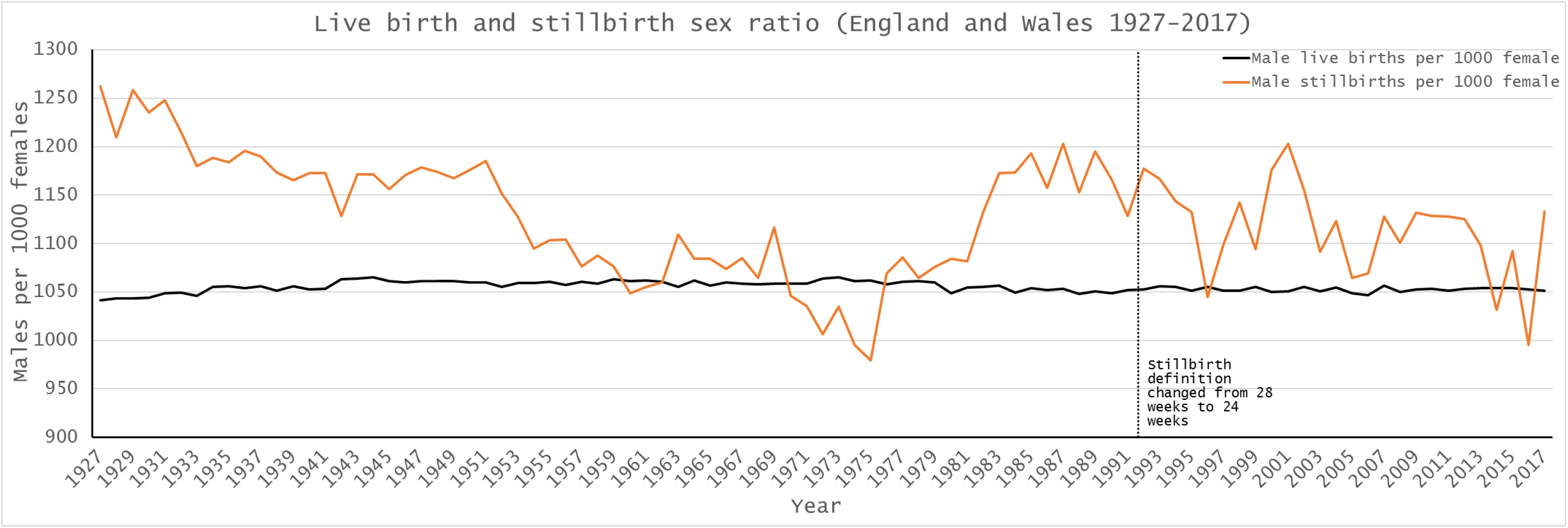
Live birth and stillbirth sex ratio in England and Wales, 1927-2017. On average 1,055 males were born per 1,000 females, and there are no years where more females are born than boys. The definition of stillbirth changed from 28 weeks of gestation to 24 weeks of gestation in 1992, and on average 1,133 boys were stillborn per 1,000 girls between 1927 and 1992, and 1,112 per 1,000 between 1993 and 2017. In the entire dataset, there are only three years where more girls were stillborn than boys (1974, 1975, 2016).

Male and female embryos differ from the very point of conception. Only males can express genes on the Y chromosome, and, prior to the completion of X chromosome inactivation, females can produce up to twice the amount of gene product for any gene encoded by the X chromosome. Male and female embryos therefore express different genes even at very early stages, varying from around 600 differentially-expressed genes in mouse blastocysts,^[16]^ to up to a third (2,921) of expressed transcripts in cow.^[17]^ Errors or delays in X chromosome inactivation, such as skewed (non-random) inactivation of the paternal or maternal copy, are likely to lead to increased fetal-maternal “differentness” and therefore preferential loss of female embryos. Indeed, such skewed inactivation has previously been implicated in recurrent miscarriage (typically defined as three or more consecutive miscarriages before 20 weeks), although evidence for this is often contradictory.^[18, 19]^ Male embryos grow faster,^[20, 21]^ and so a female embryo will be ready for implantation later than an identically-aged male, and is more likely to miss the implantation window when the endometrium is most receptive (a period of around 4 days, typically 6-8 days post-ovulation^[22]^). Male and female embryos are also metabolically distinct, as females are thought to make more use of the pentose phosphate pathway, possibly because of an additional copy of the X-linked *G6PD1* gene,^[21]^ although others have cast doubt on this idea.^[23]^ Metabolism is likely the most fundamental difference between early male and female embryos, and certainly one of the most dynamic,^[24]^ and metabolically “quiet” embryos may survive better than more active ones.^[25–27]^ In this “quiet embryo hypothesis”, metabolic signatures of the embryo are assumed to reflect viability, for instance levels of DNA damage,^[26]^ with maternal selection against metabolically-active (putatively less viable) embryos. Of course, this quest for quietness, if taken too far, would ultimately result in embryonic death, and so there must be a window of viability within which successful embryos must operate. Accepting that the endometrium acts as a biosensor to reject “unsuitable” embryos,^[6–8]^ and that there may be a “Goldilocks zone” of embryonic potential,^[28]^ females may be discriminated against from the very earliest stages of their development as early female embryos may be more different to their mother than their male counterparts. Although males have Y chromosome-specific genes that the mother does not herself possess, the X chromosome encodes more genes (846 to the 63 on the Y chromosome in Ensembl release GRCh38.p12), and prior to completion of X chromosome inactivation differential gene expression is therefore greater in females. Genes on sex chromosomes are known to regulate autosomal genes,^[23]^ and the greater number of X-linked genes in females will have a concomitantly larger effect on the number of downstream autosomal genes that are up- or down-regulated. Male and female embryos are both genetically distinct from their mother (i.e. encode paternally-derived genes), but again, because of the size of the X chromosome, females have a greater amount of paternal DNA (the X is 156Mb long, the Y just 57Mb, which may have relevance for the extent of imprinting), and a greater number of paternal genes.

A comprehensive study of the human sex ratio from conception to birth^[29]^ shows an initial large loss of male embryos in the first week or so, followed by a longer period of female-biased loss in the first trimester. As a result, the cohort sex ratio is male-biased from the end of the first trimester, and remains this way until the last few weeks of pregnancy, where male-biased stillbirth^[30]^ likely comes into play. The greatest number of female losses therefore occurs at or soon after implantation, and in the following weeks as placentation progresses. It is here that maternal-fetal contact is both established and, through placentation, extended, to reach the closest possible juxtaposition of maternal and fetal tissues and blood supplies, and this period also represents one of relatively limited fetal growth and maternal investment. It is no surprise that this should represent the “interview” period. Even once the interview is passed, the endometrium may still present a more hostile environment to female embryos, which may go some way to explaining sex differences in the male and female placenta, where female placentae are more sensitive to perturbation in the peri-conception period, and show reduced growth and a greater amount of variation in placental gene and protein expression.^[31, 32]^

### What is the extent of sex-biased loss in human pregnancy?

Whilst 10% of clinically-recognised pregnancies end in miscarriage, the true number is estimated to be much higher as many pregnancies are lost before they are identified, and up to one third of all pregnancies may end in spontaneous abortion (miscarriage).^[33–35]^ However, far higher values have been proposed.^[29, 36, 37]^ In 1975, Roberts and Lowe attempted to predict the annual number of conceptions in England and Wales in 1971,^[38]^ and suggested that up to 78% of conceptions were “lost” (unaccounted for in live birth and still birth records). Their analysis was based only on married women, hypothesised a mean frequency of coitus twice a week (one in four of which was unprotected), and did not include data for pregnancies ending in elective or therapeutic abortion. Using data from the National Surveys of Sexual Attitudes and Lifestyles (Natsal)^[39]^ it is possible to refine these calculations somewhat, and these updated calculations suggest that the assumption that around a third of all conceptions might be lost is valid (Box 1, Table 1).

**Table 1.** Theoretical prediction of the number of miscarried products of conception in England and Wales in 2012. Sexual habits are based on Natsal-3^[39]^ for women aged 16-44, and the remaining data are from ONS statistical datasets as described in the text. The predicted miscarriage rate is 43% overall, or 38% for women aged 20-39 (responsible for the majority of live births, stillbirths and abortions).

|  | Age |  |  |  |  |  |
| --- | --- | --- | --- | --- | --- | --- |
|  | Under 20 | 20-24 | 25-29 | 30-34 | 35-39 | All |
| Number of women | 1362919 | 1893629 | 1925992 | 1898383 | 1803418 | 8884341 |
| Annual acts of vaginal sex (assuming 44 per woman per year) | 59968436 | 83319676 | 84743648 | 83528852 | 79350392 | 390911004 |
| Annual acts of unprotected vaginal sex (assuming one in six is unprotected) | 9994739 | 13886613 | 14123941 | 13921475 | 13225065 | 65151834 |
| Unprotected acts occurring within 48-hour period around ovulation (i.e. 1/14) | 713910 | 991901 | 1008853 | 994391 | 944648 | 4653702 |
| Assume one in three of these results in fertilisation | 237970 | 330634 | 336284 | 331464 | 314883 | 1551234 |
| Number of live births to these women | 33815 | 132456 | 202370 | 216242 | 114797 | 699680 |
| Number of stillbirths to these women | 217 | 669 | 936 | 912 | 601 | 3118 |
| Number of elective and therapeutic abortions to these women | 30539 | 54558 | 41882 | 30353 | 18523 | 145316 |
| Estimated loss | 173399 | 142951 | 91096 | 83957 | 180962 | 672364 |
| Percentage loss | 73% | 43% | 27% | 25% | 57% | 43% |

In England and Wales, all live births and stillbirths must be registered, and there is extensive historical data available on numbers of live births and stillbirths, including sex ratios (it should be noted however that in October 1992, the Stillbirth (Definition) Act 1992 changed the gestation cut-off for stillbirths from 28 or more weeks of gestation to 24 or more weeks, and so data from 1993 onwards is not comparable to previous years). In addition to extensive live birth and stillbirth data, the requirement that all practitioners in England and Wales who perform therapeutic or elective abortions must notify the Chief Medical Officer means that abortion statistics (including number of abortions by gestation week) are available going back to the late 1960’s. The stability of the overall sex ratio at birth (Figure 1) suggests that, in England and Wales at least, there is no sex-selective abortion, and the general increase in the number of legal abortions, and the lack of maternal deaths due to complications of illegal abortions^[40]^ also suggests that there are few if any unrecorded abortions. In England and Wales between 1993 and 2017 there were 16,489,289 maternities (a pregnancy resulting in the birth of one or more children including stillbirths, of which around 1.5% resulted in multiple births); 16,656,203 live births (8,114,739 female and 8,541,464 male, with on average 1,053 males born per 1,000 females); 86,714 stillbirths (41,059 female and 45,655 male, with on average 1,112 males stillborn per 1,000 females); 4,512,024 legal elective and therapeutic abortions, and 28,269,072 conceptions (assuming that 33% of conceptions result in spontaneous abortion (miscarriage) and that the recorded live birth, stillbirth and abortion figures therefore represent 67% of total conceptions). Historically, the male bias at birth was taken to result from the production of a greater proportion of males at conception, although more recent data from *in vitro* fertilisation supports a balanced sex ratio at conception (see Orzack et al.^[29]^ for discussion), as does the simple mechanics of equal segregation of X and Y chromosomes during spermatogenesis. We can therefore reasonably conclude that the 28,269,072 conceptions comprised equal numbers of males and females. If males and females were also equally represented in the therapeutic and elective abortion dataset (i.e.the abortus sex ratio is 50:50), then 2,256,012 males and 2,256,012 females were aborted. However, more recent work^[29]^ supports a slightly female-biased cohort sex ratio during early pregnancy based on chorionic villus sampling, amniocentesis and induced abortions, and a conservative estimate of a 55:45 female:male sex ratio ≤12 weeks and 45:55 female:male ≥13 weeks might be more appropriate. In the study period, 4,044,380 therapeutic and elective abortions occurred ≤12 weeks of gestation and 467,644 occurred ≥13 weeks, with 2,435,740 girls aborted to 2,076,284 boys, for an average of 853 boys aborted per 1,000 girls (Figure 2, Supplemental table S1). Using these data, it is possible to deduce that 28,269,072 conceptions resulted in 7,014,131 spontaneous abortions (miscarriages) in England and Wales between 1993 and 2017, with on average 141,720 females and 138,845 boys lost each year, for an average miscarriage sex ratio of 980 boys per 1000 girls (Supplemental table S2). Significantly more girls are lost during pregnancy (P < 0.00001, Pearson’s χ2 test).

**Figure 2.**
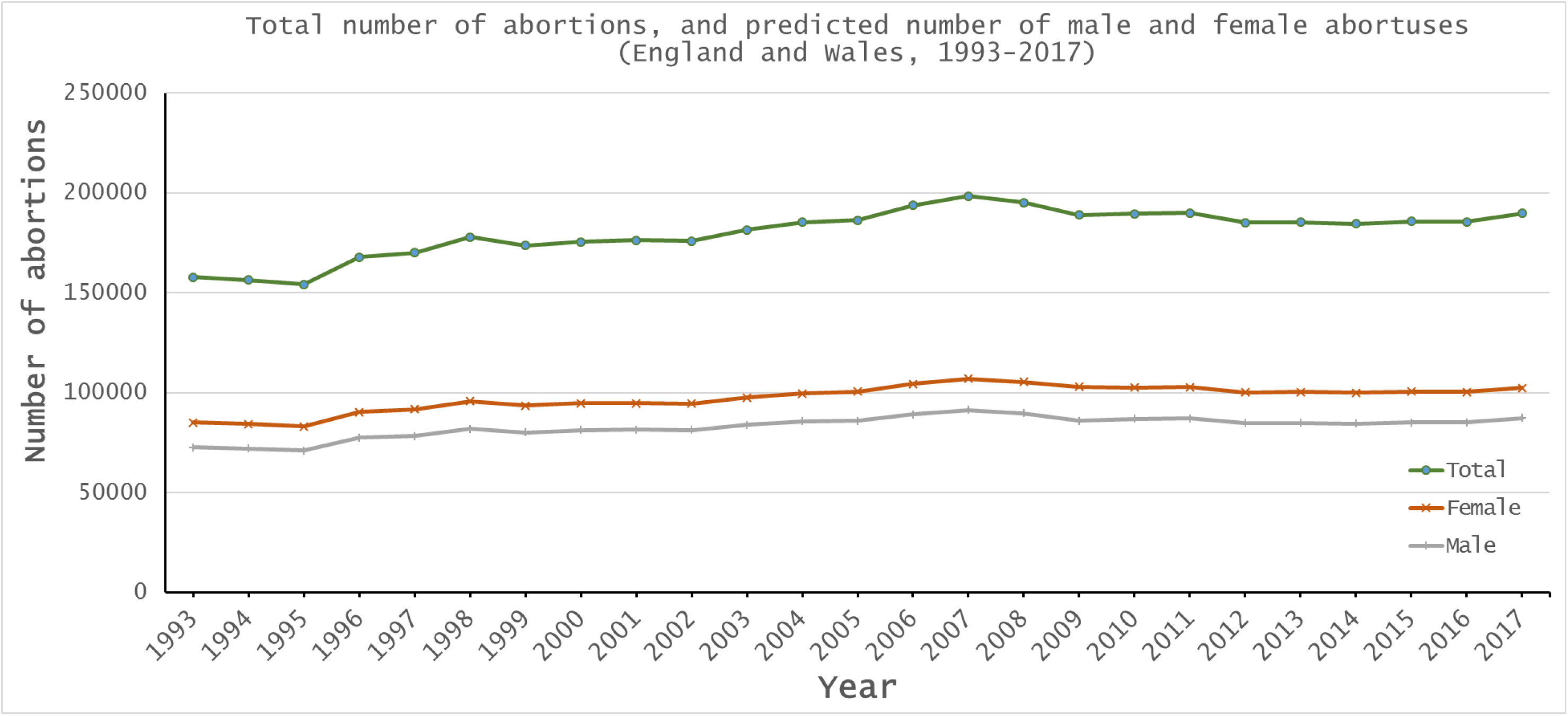
Sex ratio of therapeutic and elective abortions in England and Wales, 1993-2017, determined on the assumption that early (≤12 weeks) abortions are biased towards females (55:45) and later abortions (≥13 weeks) are biased towards males (45:55).

### Stress, miscarriage, and variation in the sex ratio at birth

The human sex ratio at birth is not stable, and, in England and Wales between 1993-2017, ranged from 1,047 boys per 1,000 girls to 1,057 boys per 1,000 girls. The predicted miscarriage sex ratio over the same period, assuming equal numbers of males and females are conceived, ranged from 956 boys per 1,000 girls in 1993 to almost parity (999 boys per 1,000 girls) in 2006 (Figure 3). The variation seen in 2006 follows the July 7^th^ 2005 terror attacks in London, and reflects the general observation that sex ratio varies following stressful events, and that parental hormone levels around conception in some ways influence the sex ratio of their offspring.^[41–43]^ Why might stress impact the human sex ratio at birth, and what exactly is changing? The human stress response is mediated by the hypothalamic-pituitary-adrenal axis, and ultimately results in the release of glucocorticoid hormones (primarily cortisol) by the adrenal cortex. Interestingly, this process also results in release of progesterone by the adrenal cortex, and leads to increased circulating levels of progesterone in serum,^[44]^ most likely because progesterone and cortisol are both cholesterol derivatives, and progesterone is a precursor in the synthesis of cortisol. Similarly, both testosterone and estradiol are cholesterol-derivatives, and so production of these hormones may also increase during the stress response. The link between cholesterol, hormones, and changes to the sex ratio at birth are hinted at in differences in ABO blood group cholesterol levels and sex ratios,^[45]^ and the link between cortisol and progesterone may also explain some seasonal variations in the human sex ratio at birth, as cortisol levels are known to vary throughout the year.^[46]^ Such a link may in future be detectable through measurement of circulating hormone levels during early pregnancy, especially if linked to early (<12 weeks) detection of fetal sex using non-invasive prenatal testing techniques (NIPT) and tracking of pregnancy outcome. If one of the factors involved in setting the threshold of acceptance for embryo implantation and peri-conception survival is the degree of differentness from the mother, then changes to hormone levels in the maternal circulation might alter the acceptance threshold, so embryos that previously would have been lost are now able to implant and survive. In particular, those female embryos produced from eggs originating in the right ovary, which would normally be rejected as too different might now survive in greater numbers.

**Figure 3.**
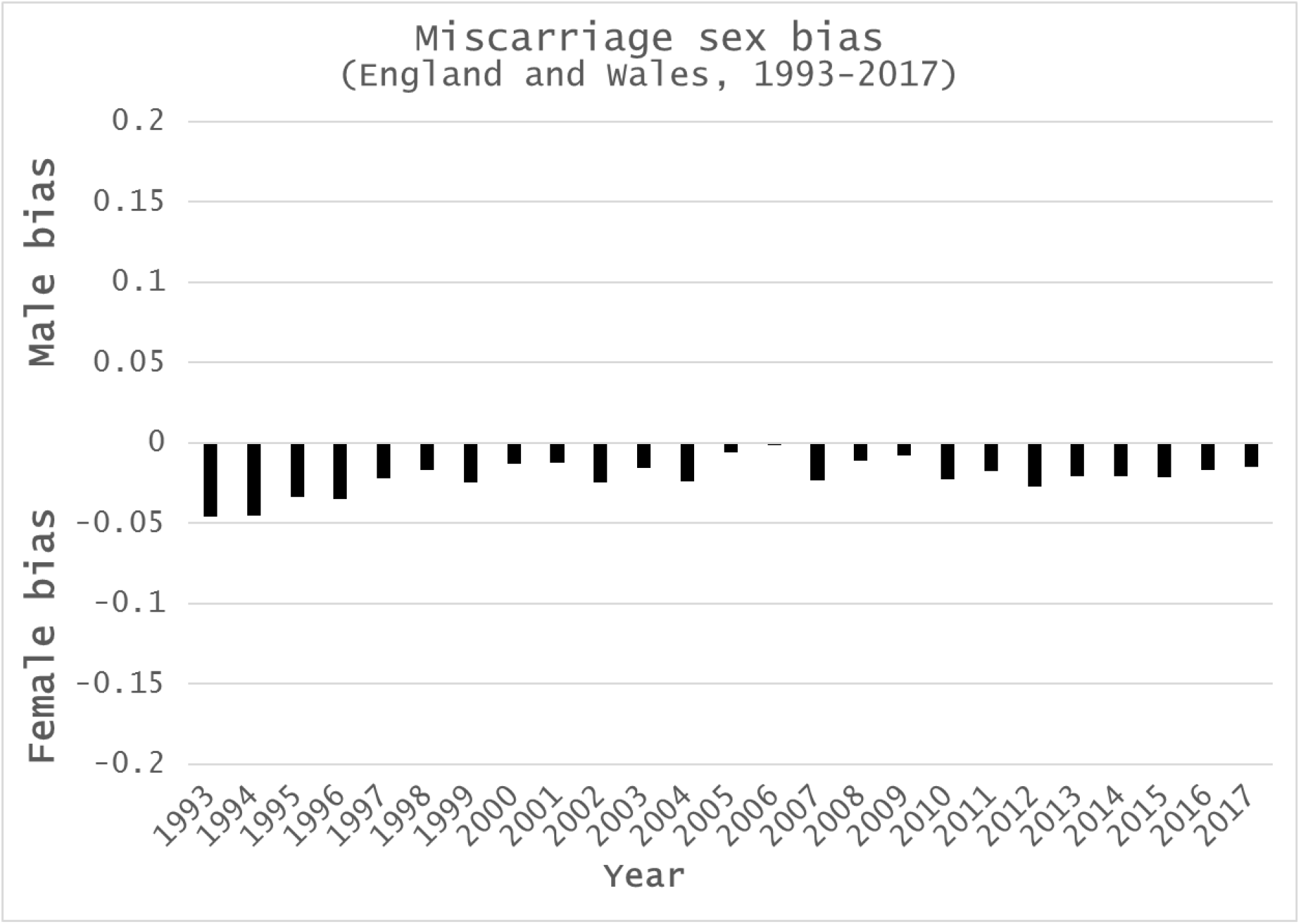
Relative miscarriage sex bias in England and Wales, 1993-2017, calculated as *1-(females miscarried/males miscarried)*, calculated if 33% of all conceptions result in miscarriage. A value of 0 indicates no bias, a positive value would show male bias, and a negative value a female bias. Miscarriages are biased towards females in every year of this 25 year dataset, with a marked decrease in the miscarriage sex ratio around 2005-2006, following the 7^th^ July 2005 terrorist bombings in London, and a smaller dip in 2009 that may be due to the 2008 financial crisis.

### Ovarian asymmetry

Humans demonstrate directional asymmetry, most obviously in the positioning of internal organs such as the heart, stomach and intestines. These asymmetries can ultimately be traced to determination of left-right axes during embryogenesis, dictated by left-biased ciliary flow and an ancient gene regulatory network involving *PITX2*, *NODAL* and *LEFTY*. Asymmetric *PITX2* expression is maintained and plays a role in subsequent organ development, and, in birds, underlies asymmetric development of the gonads, reaching its most extreme manifestation in the single (left) ovary and oviduct of many species.^[47–49]^ Among mammals, functional asymmetry of left and right ovaries has been reported from many species, including mice,^[50]^ shrews,^[51]^ gerbils,^[52]^ viscachia,^[53, 54]^ bats,^[55]^ and waterbuck.^[56]^ Although human gonadal asymmetry is perhaps most apparent in males, where the right testis is larger, the inherent directional asymmetry of vertebrate embryos demonstrates that the human left and right ovary are not equal from the earliest stages of development. In adults, this asymmetry manifests itself in anatomical relations (the left ovary lies adjacent to the sigmoid colon, the right nearer the appendix), venous drainage (the left ovary drains into the left renal vein, the right into the inferior vena cava^[57, 58]^), and function. The right ovary may ovulate more frequently and favour pregnancy,^[59–63]^ and this elevated function possibly leaves the right ovary more susceptible to ovarian cancer,^[64, 65]^ cystic ovarian endometriosis,^[66]^ and ruptured corpus luteum.^[57, 67–69]^ Ectopic pregnancy may also be more common on the right,^[70–72]^ and gonadal tissues are unevenly distributed in true hermaphrodites, with ovaries more common on the left, and testes/ovotestes more common on the right.^[73–75]^ The data on functional asymmetry of human ovaries and possible differential susceptibility to disease are noisy with generally small effects, however, the consistent trends, coupled with developmental and anatomical asymmetries, tells us that we should not consider left and right ovaries as equals. What implications might ovarian asymmetry have for human reproduction? There are hints in the literature that the right ovary might ovulate more, and favour pregnancy,^[59–63]^ but one possibility that is generally neglected is that differences between left and right ovaries might lead to variation in the human sex ratio at birth. It has long been recognised that more males are born than females in many populations, despite greater susceptibility of boys to stillbirth,^[30]^ (Figure 1). Whilst the sex ratio at birth is typically stable, and for the most part biased towards males, there is variation across populations, only some of which is likely due to sex-specific elective abortion.^[76]^ The remaining variation can seemingly be explained by demography, with those of African origin having lower sex ratios^[77, 78]^ (even becoming female-biased in some cases^[77]^), latitude,^[79]^ and seasonality.^[80–82]^ Perhaps most interestingly, the sex ratio at birth can be perturbed by stressful events, such as the 1995 Kobe earthquake;^[83]^ famine;^[84]^ war;^[85–87]^ terrorist attacks;^[88–90]^ historic royal events;^[91]^ the Superbowl;^[92]^ and economic stress.^[93, 94]^ Such seasonal variation, coupled with the effects of stress, is evidence for hormonal influences on the human sex ratio at birth, and, indeed, it has previously been suggested that hormonal concentrations in parents around conception can alter sex ratios.^[41, 95]^ Once asymmetry of left and right ovaries is accepted, these influences may become easier to explain. The right and left ovaries differ in their venous drainage, and as a result, pressure in the right ovarian vein is higher than the left.^[57, 58]^ The right ovary therefore drains more slowly than the left, and so we can expect hormones to accumulate differentially, resulting in higher concentrations of estradiol, testosterone, progesterone, follicle-stimulating hormone (FSH), luteinizing hormone (LH), and cortisol on this side. These elevated concentrations are maintained, and perhaps boosted, by counter current exchange between ovarian veins and arteries.^[96]^ Prior to ovulation, a human oocyte is bathed in approximately 5ml of follicular fluid, containing estradiol and progesterone in concentrations roughly 500x and 1000x higher respectively than in serum,^[97–99]^ and the freedom of steroid hormones to move across membranes means that the oocyte will equilibrate to follicular fluid conditions prior to ovulation. Oocytes from the right and left ovaries will therefore experience different environments as folliculogenesis progresses, with levels of estradiol, progesterone and others far in excess of maternal serum levels, and they will carry these hormones with them in their cytoplasm (and that of their companion cells) as they are released. At the same time, several millilitres of asymmetric follicular fluid is released, producing different microenvironments in the left and right fallopian tubes as the egg begins its journey towards fertilisation.

### Ovarian asymmetry and human reproduction

Human sperm are attracted to follicular fluid.^[100, 101]^ More specifically, a subset of sperm cells undergo capacitation, and as a result demonstrate chemotaxis,^[102]^ with low levels of progesterone known to act as a chemoattractant.^[103–105]^ There is some debate regarding the role that follicular fluid plays *in vivo*, with evidence from animal studies suggesting ≤1% of the follicular fluid released along with the egg actually enters the fallopian tube,^[106–109]^ equating to ≤50μl in humans. Others have suggested that follicular fluid is the major fluid constituent in fallopian tubes immediately post-ovulation, as the egg is carried into the tube via a wave of fluid.^[110]^ Follicular fluid that does not directly enter the fallopian tube is released into the peritoneal cavity immediately adjacent to the ovary, and the ruptured follicle may continue to secrete follicular fluid for a short time following ovulation (the developing follicle may also secrete hormones into the peritoneal cavity for a period *prior* to ovulation^[111]^). There is therefore a pool of hormone-enriched fluid adjacent to the fimbria at the end of the fallopian tube before and after ovulation, and this may be drawn into the tube by ciliary flow^[112, 113]^ or enter adjacent blood vessels. Since the composition of fallopian tube fluid differs between left and right sides, and since sperm respond to progesterone concentrations in the picomolar range,^[102, 114]^ these differences may have implications for sperm chemotaxis. Given that the follicular microenvironment is virtually saturated with progesterone, the oocyte and its associated cumulus cells are likely equilibrated to follicular fluid conditions, and the oocyte-cumulus complex itself is also a source of chemoattractants, including progesterone.^[109, 114, 115]^ We therefore have a situation where the chemoattractant concentrations within oocyte-cumulus complexes differs between left and right sides, and where the fallopian tube fluid that bathes these complexes also differs.

With this in mind, there are several ways that ovarian asymmetry (as demonstrated by variation in hormonal concentrations in follicular fluid from left and right ovaries) might influence human fertility. Firstly, progesterone can inhibit ciliary beating in the fallopian tube,^[116, 117]^ and so egg motility may differ between the left and right tubes, with extended migration times reducing the possibility of “healthy” sperm reaching the egg. Secondly, different hormone/chemoattractant concentrations on the left and right may affect sperm chemotaxis, either positively, through improved or earlier attraction of sperm, or negatively, through saturation of receptors. Saturated receptors can no longer detect increases in chemoattractant concentration, rendering chemotaxis impossible, and studies of human sperm cell responses to progesterone have indeed shown that high concentrations are ineffective, and that sperm more readily respond to concentrations in the picomolar range.^[102, 114]^ These effects may operate over both long (from the sperm reservoir to the oocyte-cumulus complex) or short (within the oocyte-cumulus complex) distances. Similarly, X and Y chromosome-bearing sperm may respond differently to signals from the right and left. There has been much debate over morphological or behavioural differences between X and Y chromosome-bearing sperm, with some suggesting that those carrying the smaller Y chromosome may have a smaller head size and swim faster (and potentially further) than those bearing the larger X chromosome.^[118, 119]^ The difference in DNA content between X and Y chromosome-bearing sperm is roughly 3%, and, although small, it does seem likely that this has at least some influence on size and/or shape of the sperm head.^[21, 120–122]^ Sperm carrying X or Y chromosomes may therefore have either variable numbers of chemoreceptors such as CatSper^[123, 124]^ and hOR17-4,^[125]^ or these receptors may be distributed differently across the sperm head. A higher number of receptors on larger X chromosome-bearing sperm might improve sensitivity, whilst a lower number on those carrying a Y may make them more easily saturated. Variability in receptor distribution across larger or smaller sperm heads might also improve directionality, or simply improve sensitivity by widening the detection window. Most importantly, the different intrinsic hormone concentrations of eggs originating in the left and right ovaries might impact embryonic implantation, placentation, and post-implantation survival and especially greater survival of female embryos during times of stress when the maternal acceptance threshold shifts in their favour.

The literature concerning the human sex ratio at birth is very male-centric. Discussion of variation in sex ratio, and especially declines, typically assumes that this is the result of a greater loss of males (the “fragile male” idea^[126–128]^), possibly because most male losses occur later in development, and so are more visible. A similar result can of course also be explained by more females surviving than would usually be the case. ^[129]^ Such increased survival can affect the sex ratio at birth in several ways, most obviously with a greater number of female live births, but also by impacting subsequent pregnancies. In those actively trying to conceive, a female embryo lost early (at or soon after implantation) might have been replaced by a male in a subsequent cycle, but survival of the female embryo removes that mother from the pool of potential reproducers for the duration of the pregnancy, and sometime beyond. Reproductive behaviour may also be important, particularly in terms of stopping rules,^[130]^ where couples desiring a child of a specific sex stop reproducing once this is achieved, or where couples might wish for a child of each sex, and continue reproducing until this is achieved. In conditions which favour survival of females, those seeking a girl might therefore stop reproducing after one pregnancy, and those who seek a boy and a girl would stop if they already have a boy. Conversely, those seeking a boy who already have one girl might continue to reproduce after having another, and, if the stressful conditions endure, may continue to have girls. However, it must be kept in mind that changes to the human sex ratio at birth are typically small, varying only between a low of 51.02:48.98% male:female in 1927 to a high of 51.58:48.42% male:female in 1973, based on live birth data for England and Wales from 1927-2017. In the 62,454,461 live births recorded during this period, only 1,685,618 more boys than girls were born. If slightly more boys were conceived (i.e. if the primary sex ratio at conception was not exactly equal), then this miscarriage sex bias would disappear – althoughy we would then need a mechanism which would favour greater conception of boys. Whilst ovarian asymmetry and a threshold dichotomy of implantation and placentation success predicated upon uterine biosensing may not account for all of the observed variation in miscarriage sex, it does represent a novel mechanism by which we can explain existing data, such as the influence of maternal hormones around the time of conception, and the impact of stressful events and seasonality on the human sex ratio at birth.

## Conclusions

Ovarian asymmetry is a neglected aspect of reproductive biology. It is rare to find a scientific publication dealing with embryos produced by assisted reproductive technology or the composition of follicular fluid that identifies from which ovary the study materials were sourced. Similarly, studies of implantation rarely, if ever, identify the sex of embryos concerned. Research in adult humans, or using animals, is required to report the participant sex, and it is perhaps time for greater awareness of embryonic sex and ovary-of-origin in the study of early human development. A greater appreciation of ovarian asymmetry may also be necessary for explaining variation within and between women (Box 2). Similarly, consideration of testicular asymmetry, and that of the uterus and fallopian tubes, may also be overdue, as early developmental asymmetries likely impact all of these structures.

More girls are lost during pregnancy than boys, and as a result the human sex ratio at birth is biased towards males. Male and female embryos are not equal from the very moment of conception, and it should be no surprise that these differences might influence some of the most important aspects of mammalian development, such as implantation and placentation. The greater genetic and metabolic “differentness” of female embryos, at least prior to the development of functional gonads, may count against them in the threshold dichotomy^[10]^ of acceptance or rejection by maternal tissues. Such discrimination may ultimately work in their favour however, if it follows a pattern similar to that in ‘reverse’ imprinting,^[131, 132]^ where expression of maternal alleles might be favoured if elevated gene expression increases the possibility of spontaneous abortion, but leads to an increase in robustness (increased growth and pre- and post-natal survival) of survivors. If losses occur early in pregnancy, minimal resources have been invested and cost to the mother is limited. Boys exhibit greater infant mortality than girls^[133]^ (and higher stillbirth rates) and so it may be that the greater loss of girls earlier in pregnancy actually explains their later robustness.

## Acknowledgements

I would like to thank Tom Brekke, Daniel Brison, Henry Leese, John Aplin, and David Steinsaltz for comments and advice on the manuscript, and Gudrun Rieck for general discussion. Comments from Professor Bill James and two anonymous reviewers greatly improved the final manuscript.

### Box 1. Calculating the number of missing conceptions

Natsal-3^[39]^ suggests that women aged 16-44 in the survey period (6^th^ September 2010 – 31^st^ August 2012) had on average 4.9 occasions of sexual intercourse (defined as vaginal, oral or anal intercourse) in the preceding 4 weeks, of which 69% included vaginal sex (defined as a man’s penis in a woman’s vagina). The frequency of sexual intercourse for this age group is likely nearer 1.2 occasions per week, with 0.85 instances of vaginal sex per week. The annual frequency (i.e. for 52 weeks) of vaginal sex for women aged 16-44 in the survey period was therefore 44.2, not the 104 previously used by Roberts and Lowe.^[38]^ The proportion of unprotected acts of coitus during the survey period is also lower than the 25% estimate of Roberts and Lowe, and is likely nearer 5-7% for women aged 16-44,^[134]^ increasing to around 10% if less effective methods of contraception are included, or to 1/6 if some consideration is given to those trying to conceive or who were already pregnant. The number of unprotected instances of vaginal sex per woman per year is therefore around 7, and, of these, 1/14 will occur within 48 hours of ovulation. Given a fertilisation rate of around 60% in *in vitro* fertilisation,^[135]^ where sperm quality and quantity is likely higher than that of a “normal” ejaculate at the point of fertilisation *in vivo*, a fertilisation rate of one in three seems reasonable. The Office for National Statistics mid-year population estimate for mid-2012 predicted that there were 8,884,341 women between the ages of 16-39 in England and Wales, and so the estimated the number of “missing” conceptions (i.e. those not accounted for in the relevant live birth, stillbirth, and legal therapeutic and elective abortion statistics) for women aged 16-39 in 2012 was 43%. For women aged 20-39 (responsible for 91% of all live births, 88% of stillbirths and 78% of abortions), the average rate of loss was 38% (Table 1). Given the inherent uncertainty in these calculations, an estimate that around a third of all conceptions are lost seems reasonable.

### Box 2. Quantifying ovarian asymmetry

We know that the right and left ovaries are not equal. They originate on different sides of an asymmetric embryo, and lie in different sides in a directionally asymmetric adult. Differences in venous relations suggest that rates of drainage will vary, and we might therefore expect that levels of various hormones and metabolites might also vary. What is needed now is direct measurement of these variations. Whilst this may seem a relatively simple experiment, requiring only collection of follicular fluid from the left and right ovaries of a large number of women during assisted-reproduction, it is complicated by the fact that every ovulation changes the ovary, forever, converting a primordial follicle into a fibrous, scar-like corpus albicans. However, ovaries do not endlessly accumulate corpora albicantia, and those of premenopausal women undergo a process of fibroblastic replacement, ultimately forming a new section of ovarian connective tissue (stroma). Given different patterns of right/left ovulation between women, and with variable gaps related to childbirth or contraception, it becomes clear that we should be very careful when comparing ovaries between even age-matched women. How then might we quantify ovarian asymmetry in the face of differential patterns and numbers of ovulations? The simplest approach seems to be to go earlier, and to investigate inherent ovarian asymmetry before the onset of puberty, through measurement of levels of hormones and metabolites in the left and right ovaries with respect to circulating levels.

## Supplemental information

### Supplemental methods

Data for numbers of maternities, live births and stillbirths, including numbers of males and females, were collected from the Office for National Statistics (https://www.ons.gov.uk/) ‘Review of the Registrar General on births and patterns of family building in England and Wales’, Series FM1 (numbers 22-37, covering 1993-2008), the ‘Characteristics of Birth 2, England and Wales’ dataset (2009-2013), the ‘Birth characteristics dataset’ (2014-2016), and the ‘Summary of key birth statistics, 1838 to 2017’. Data on numbers of legal abortions from 2011-2017 were obtained from the Department of Health and Social Care (DHSC) ‘Abortion statistics, England and Wales’ collection (https://www.gov.uk/government/collections/abortion-statistics-for-england-and-wales), and for 1993-2010 from the UK Government Web Archive (http://www.nationalarchives.gov.uk/webarchive/). Numbers of stillbirths by age of mother for 2012 were obtained from the from the ‘Child Mortality Statistics 2012’ dataset (https://www.ons.gov.uk/peoplepopulationandcommunity/birthsdeathsandmarriages/deaths/datasets/childmortalitystatisticschildhoodinfantandperinatalchildhoodinfantandperinatalmortalityinenglandandwales), and England and Wales population data were obtained from the ‘MYE2: Population Estimates by single year of age and sex for local authorities in the UK, mid-2012’ (https://www.ons.gov.uk/peoplepopulationandcommunity/populationandmigration/populationestimates/datasets/populationestimatesforukenglandandwalesscotlandandnorthernireland).

Gestation week data is variable across the abortion dataset, and so statistics were pooled into abortions occurring either ≤12 weeks or ≥13 weeks. A comprehensive study of the human sex ratio from conception to birth^[29]^ supports a female-biased cohort sex ratio during early pregnancy based on chorionic villus sampling, amniocentesis and induced abortions, and I have therefore chosen a conservative estimate of a 55:45 female:male sex ratio ≤12 weeks and 45:55 female:male ≥13 weeks. Using these values, I calculated the number of male and female abortuses each year.

Adding together the total number of live births, stillbirths and legal abortions provided the number of pregnancies, and accepting that these represent 67% of actual conceptions (i.e. the probability that a conception resulted in miscarriage was 33%) determined the relevant number of conceptions. If the primary sex ratio is equal, then equal numbers of males and females are conceived, and subtraction of the known numbers of live and stillborn males and females, and the predicted male and female abortuses left the number of products of conception lost to miscarriage.

Statistical significance of deviation of calculated numbers of miscarried males and females from expected numbers (males and females are equally susceptible to miscarriage) was assessed using Pearson’s χ^2^ test. All calculations were rounded to the nearest whole number to reflect the impossibility of conceiving a fraction of a person, and so in some cases annual totals are not the sum of their constituent parts. It also goes without saying that the ratios presented here address only a narrow range of biological sex, not gender, and are predicated on the simplistic assumption that XX = female and XY = male.

---

**Supplemental table S1.** Live births, stillbirths and abortions in England and Wales, 1993-2017. Numbers of males and females for live births and stillbirths reflect classifications as recorded on the relevant birth registers. Abortus sex is calculated from the total number of therapeutic and elective abortions on the assumption that the sex ratio ≤12 weeks of gestation is 55:45 in favour of females, and 45:55 in favour of males from ≥13 weeks of gestation. Total or average values are provided in the bottom rows.

| Year | Live births |  |  |  | Stillbirths |  |  |  | Abortions |  |  |  | Total (live births + stillbirths + abortions) |
| --- | --- | --- | --- | --- | --- | --- | --- | --- | --- | --- | --- | --- | --- |
|  | Total | Female | Male | Male live births per 1,000 female live births | Total | Female | Male | Male stillbirths per 1000 female stillbirths | Total | Female | Male | Male abortuses per 1000 female abortuses |  |
| 1993 | 673467 | 327632 | 345835 | 1056 | 3855 | 1779 | 2076 | 1167 | 157846 | 85072 | 72775 | 855 | 835168 |
| 1994 | 664726 | 323405 | 341321 | 1055 | 3813 | 1779 | 2034 | 1143 | 156539 | 84363 | 72176 | 856 | 825078 |
| 1995 | 648138 | 315950 | 332188 | 1051 | 3600 | 1688 | 1912 | 1133 | 154315 | 83211 | 71104 | 854 | 806053 |
| 1996 | 649485 | 315995 | 333490 | 1055 | 3539 | 1731 | 1808 | 1044 | 167916 | 90444 | 77472 | 857 | 820940 |
| 1997 | 642093 | 313021 | 329072 | 1051 | 3439 | 1638 | 1801 | 1100 | 170145 | 91732 | 78413 | 855 | 815677 |
| 1998 | 635901 | 309998 | 325903 | 1051 | 3417 | 1595 | 1822 | 1142 | 177871 | 95875 | 81996 | 855 | 817189 |
| 1999 | 621872 | 302617 | 319255 | 1055 | 3305 | 1578 | 1727 | 1094 | 173701 | 93634 | 80067 | 855 | 798878 |
| 2000 | 604441 | 294816 | 309625 | 1050 | 3203 | 1472 | 1731 | 1176 | 175542 | 94485 | 81057 | 858 | 783186 |
| 2001 | 594634 | 289999 | 304635 | 1050 | 3159 | 1434 | 1725 | 1203 | 176364 | 94851 | 81513 | 859 | 774157 |
| 2002 | 596122 | 290059 | 306063 | 1055 | 3372 | 1565 | 1807 | 1155 | 175932 | 94542 | 81390 | 861 | 775426 |
| 2003 | 621469 | 303041 | 318428 | 1051 | 3585 | 1711 | 1874 | 1091 | 181582 | 97557 | 84025 | 861 | 806636 |
| 2004 | 639721 | 311381 | 328340 | 1054 | 3686 | 1736 | 1950 | 1123 | 185415 | 99690 | 85725 | 860 | 828822 |
| 2005 | 645835 | 315235 | 330600 | 1049 | 3483 | 1687 | 1796 | 1065 | 186416 | 100535 | 85881 | 854 | 835734 |
| 2006 | 669601 | 327172 | 342429 | 1047 | 3602 | 1741 | 1861 | 1069 | 193737 | 104469 | 89268 | 854 | 866940 |
| 2007 | 690013 | 335525 | 354488 | 1057 | 3598 | 1691 | 1907 | 1128 | 198499 | 107139 | 91360 | 853 | 892110 |
| 2008 | 708711 | 345748 | 362963 | 1050 | 3617 | 1722 | 1895 | 1100 | 195296 | 105514 | 89782 | 851 | 907624 |
| 2009 | 706248 | 344113 | 362135 | 1052 | 3688 | 1730 | 1958 | 1132 | 189100 | 103114 | 85986 | 834 | 899036 |
| 2010 | 723165 | 352199 | 370966 | 1053 | 3714 | 1745 | 1969 | 1128 | 189574 | 102582 | 86992 | 848 | 916453 |
| 2011 | 723913 | 352939 | 370974 | 1051 | 3811 | 1791 | 2020 | 1128 | 189931 | 102787 | 87144 | 848 | 917655 |
| 2012 | 729674 | 355328 | 374346 | 1054 | 3558 | 1674 | 1884 | 1125 | 185122 | 100150 | 84972 | 848 | 918354 |
| 2013 | 698512 | 340129 | 358383 | 1054 | 3284 | 1565 | 1719 | 1098 | 185331 | 100360 | 84971 | 847 | 887127 |
| 2014 | 695233 | 338461 | 356772 | 1054 | 3254 | 1602 | 1652 | 1031 | 184571 | 99998 | 84573 | 846 | 883058 |
| 2015 | 697852 | 339716 | 358136 | 1054 | 3147 | 1498 | 1649 | 1092 | 185824 | 100649 | 85175 | 846 | 886823 |
| 2016 | 696271 | 339225 | 357046 | 1053 | 3112 | 1560 | 1552 | 995 | 185596 | 100501 | 85095 | 847 | 884979 |
| 2017 | 679106 | 331035 | 348071 | 1051 | 2873 | 1347 | 1526 | 1133 | 189859 | 102488 | 87371 | 853 | 871838 |
| Total: | 16656203 | 8114739 | 8541464 | - | 86714 | 41059 | 45655 | - | 4512024 | 2435740 | 2076284 | - | 21254941 |
| Average: | 666248 | 324590 | 341659 | 1053 | 3469 | 1642 | 1826 | 1112 | 180481 | 97430 | 83051 | 853 | 850198 |

**Supplemental table S2.** Predicted spontaneous abortions (miscarriages) in England and Wales, 1993-2017. The number of conceptions is calculated on the assumption that the sum of live births, stillbirths and elective and therapeutic abortions represents 67% of all conceptions (i.e. 33% of conception are lost). The sex ratio at conception is equal, and so the number of miscarriages can be calculated from the number of males or females conceived and the number accounted for in live birth, stillbirth or abortion statistics. Total or average values are provided in the bottom rows.

|  | Number of conceptions |  |  | Number of conceptions accounted for in live birth stillbirth and abortion data |  |  | Number of miscarriages |  |  |  |
| --- | --- | --- | --- | --- | --- | --- | --- | --- | --- | --- |
|  | Total | Female | Male | Total | Female | Male | Total | Female | Male | Male miscarriages per 1000 female miscarriages |
| **1993** | 1110773 | 555387 | 555387 | 835168 | 414483 | 420686 | 275605 | 140904 | 134701 | 956 |
| **1994** | 1097354 | 548677 | 548677 | 825078 | 409547 | 415531 | 272276 | 139130 | 133146 | 957 |
| **1995** | 1072050 | 536025 | 536025 | 806053 | 400849 | 405204 | 265997 | 135176 | 130822 | 968 |
| **1996** | 1091850 | 545925 | 545925 | 820940 | 408170 | 412770 | 270910 | 137755 | 133155 | 967 |
| **1997** | 1084850 | 542425 | 542425 | 815677 | 406391 | 409286 | 269173 | 136034 | 133139 | 979 |
| **1998** | 1086861 | 543431 | 543431 | 817189 | 407468 | 409721 | 269672 | 135963 | 133709 | 983 |
| **1999** | 1062508 | 531254 | 531254 | 798878 | 397829 | 401049 | 263630 | 133425 | 130205 | 976 |
| **2000** | 1041637 | 520819 | 520819 | 783186 | 390773 | 392413 | 258451 | 130046 | 128405 | 987 |
| **2001** | 1029629 | 514814 | 514814 | 774157 | 386284 | 387873 | 255472 | 128531 | 126941 | 988 |
| **2002** | 1031317 | 515658 | 515658 | 775426 | 386166 | 389260 | 255891 | 129492 | 126399 | 976 |
| **2003** | 1072826 | 536413 | 536413 | 806636 | 402309 | 404327 | 266190 | 134104 | 132086 | 985 |
| **2004** | 1102333 | 551167 | 551167 | 828822 | 412807 | 416015 | 273511 | 138360 | 135151 | 977 |
| **2005** | 1111526 | 555763 | 555763 | 835734 | 417457 | 418277 | 275792 | 138306 | 137486 | 994 |
| **2006** | 1153030 | 576515 | 576515 | 866940 | 433382 | 433558 | 286090 | 143133 | 142957 | 999 |
| **2007** | 1186506 | 593253 | 593253 | 892110 | 444355 | 447755 | 294396 | 148898 | 145498 | 977 |
| **2008** | 1207140 | 603570 | 603570 | 907624 | 452984 | 454640 | 299516 | 150586 | 148930 | 989 |
| **2009** | 1195718 | 597859 | 597859 | 899036 | 448957 | 450079 | 296682 | 148902 | 147780 | 992 |
| **2010** | 1218882 | 609441 | 609441 | 916453 | 456526 | 459927 | 302429 | 152915 | 149514 | 978 |
| **2011** | 1220481 | 610241 | 610241 | 917655 | 457517 | 460138 | 302826 | 152724 | 150102 | 983 |
| **2012** | 1221411 | 610705 | 610705 | 918354 | 457152 | 461202 | 303057 | 153554 | 149503 | 974 |
| **2013** | 1179879 | 589939 | 589939 | 887127 | 442054 | 445073 | 292752 | 147885 | 144867 | 980 |
| **2014** | 1174467 | 587234 | 587234 | 883058 | 440061 | 442997 | 291409 | 147172 | 144237 | 980 |
| **2015** | 1179475 | 589737 | 589737 | 886823 | 441863 | 444960 | 292652 | 147874 | 144777 | 979 |
| **2016** | 1177022 | 588511 | 588511 | 884979 | 441286 | 443693 | 292043 | 147225 | 144818 | 984 |
| **2017** | 1159545 | 579772 | 579772 | 871838 | 434870 | 436968 | 287707 | 144903 | 142804 | 986 |
| **Total:** | **28269072** | **14134536** | **14134536** | **21254941** | **10591538** | **10663403** | **7014131** | **3542998** | **3471133** | **-** |
| **Average:** | **1130763** | **565381** | **565381** | **850198** | **423662** | **426536** | **280565** | **141720** | **138845** | **980** |

